# Intracellular calcium elevations drive the nucleation of FIP200- and ATG13-containing pre-autophagosomal structures that become omegasomes

**DOI:** 10.1101/2022.11.02.514842

**Authors:** Matthew Smith, Priya Schoenfelder, Maria Manifava, Hannah Polson, Sharon Tooze, H Llewelyn Roderick, Nicholas T. Ktistakis

## Abstract

Ca2+ modulates autophagy at multiple steps including the induction and maturation of autophagosomes, but the magnitude and spatiotemporal properties of this calcium signal and its ultimate effect on the autophagic machinery are unclear. Focusing on the induction step leading to omegasome formation, we report that low but sustained elevations in cytosolic calcium levels induce omegasome formation but treatments that only transiently elevate calcium do not. The calcium-induced structures are early intermediates that mature into omegasomes but do not constitute full autophagosomes because they are partially devoid of late autophagy proteins ATG16 and LC3. In addition to omegasomes, all four components of the ULK complex (ULK1, FIP200, ATG13, ATG101) respond to calcium modulation: they translocate to early autophagy puncta in complete medium upon calcium elevation, and are inhibited from translocation during starvation by calcium chelation with BAPTA-2 AM. The principal early step affected by calcium lies downstream of mTORC1 inactivation and upstream of VPS34 activation, coinciding biochemically with phosphorylation of ATG13 at serine 318, which is known to require ULK1 activity. However, although the calcium-mediated step requires ATG9, FIP200 and ATG13, it does not require ULK1/2, suggesting that calcium does not directly regulate ULK1 activity but rather it regulates the mechanism by which the ULK complex components ATG13 and FIP200, together with ATG9, nucleate pre-autophagosomal precursors. This calcium-induced nucleation is sufficient to drive autophagy induction up to the omegasome step, but not beyond it.

## Introduction

Macroautophagy (hereafter autophagy) is a highly conserved, intracellular catabolic mechanism that functions through the sequestration of cytoplasmic cargo into double-membraned vesicles, termed autophagosomes, either selectively or non-selectively. Autophagosomes ultimately fuse with lysosomes, resulting in degradation of autophagic cargo[1].

The machinery involved in autophagy requires numerous complexes to co-operate systematically for its function. The earliest complex required for autophagy is the unc-51-like kinase (ULK) 1 complex, which consists of the serine-threonine kinase ULK1 and the scaffold proteins focal adhesion kinase family interacting protein of 200 kDa (FIP200), autophagy-related protein (ATG) 13 [2] and ATG101 [3]. Whereas the ULK complex is widely believed to be essential for amino acid starvation-induced autophagy, recent evidence suggests that ULK-independent autophagy pathways may exist, especially in response to specific autophagic stimuli such as glucose starvation or mitochondrial damage [4-7].

Mechanistically, upon autophagy activation, the ULK complex translocates to the site of autophagy induction (known as the phagophore) to which it recruits a second initiation complex, the vacuolar sorting protein 34 (VPS34) complex, which synthesises phosphatidylinositol 3-phosphate (PI3P) through the activity of the lipid kinase VPS34. PI3P itself can then recruit PI3P-effector proteins, such as double FYVE containing protein 1 (DFCP1) and WD repeat domain phosphoinositide-interacting proteins (WIPIs) at a site termed the omegasome [8, 9], within which the phagophore expands and closes. ATG9 is also required for autophagy initiation. This transmembrane protein shuttles from various cellular membranes to interact with the phagophore and omegasomes [10]. The final complex that is recruited is the ATG16L1–ATG5–ATG12 conjugation machinery, which acts to catalyse the conjugation of ATG8-family proteins (microtubule-associated protein 1 light chain 3 [LC3], or GABA type A receptor associated protein [GABARAP]) to phosphatidylethanolamine, a key requirement in autophagosome closure and fusion with lysosomes [11, 12].

Calcium (Ca^2+^) is a key cellular messenger that regulates a vast array of cellular functions. Ca^2+^ performs its function through distinct spatiotemporal signals that are subsequently decrypted by the cell, resulting in the activation of specific signalling cascades [13]. The source of this Ca^2+^ can be extracellular, or is located in intracellular Ca^2+^ stores including the ER and acidic organelles [14].

The role of Ca^2+^ in regulating autophagy has been complicated to discern, with various studies suggesting either an inhibitory or activating effect [15]. A majority of the evidence supporting a suppressive effect comes from the finding that inhibition of ER-localised Ca^2+^ channels leads to an increase in autophagy [16-19]. However, there appears to be a larger body of evidence supporting a pro-autophagic effect of Ca^2+^. From these reports, the most common downstream effector pathway that has been suggested to sense and be modulated by cytosolic Ca^2+^ is the Ca^2+^-calmodulin-dependent protein kinase kinase-β/ AMP-activated protein kinase/ mammalian target of rapamycin (CaMKKβ/ AMPK/ mTOR) cascade [20-24], although an AMPK-independent mechanism has also been proposed [25]. Additionally, PKCθ [26], ERK [27] and Beclin1 [28] have also been proposed as alternate targets. Further support for an activating role of Ca^2+^ in autophagy is provided by evidence that chelating cytosolic Ca^2+^ inhibits autophagic induction, not only in response to Ca^2+^ mobilisation, but also when cells are stimulated through classical starvation-induced signals [22, 29, 30].

In addition to the debate on the effect of Ca^2+^ in autophagy, the identity of its source has also been of much debate. To date, Transient Receptor Potential Channel Mucolipin (TRPML) 1 [31-36], TRPML3 [37-39], two-pore channel (TPC) 2 [40, 41], L-type calcium channels [42], T-type calcium channels [43], mitochondrial calcium uniporter [44, 45] and the inositol trisphosphate receptor (IP_3_R) [27, 46] have all been proposed as the channel(s) responsible for generating the Ca^2+^ signal underlying the induction of autophagy.

Despite an increasing number of reports implicating Ca^2+^ in autophagy, the conditions under which it elicits an inhibitory or an activating effect are not clear. Recent studies from the Zhang lab have shed light on this problem, suggesting that ER Ca^2+^ release and generation of a Ca2+ signal localised to the ER membrane is involved in the formation of FIP200-dependent pre-autophagosomal structures through the involvement of the VMP1 and EI24 proteins [47, 48]. In the work presented in this manuscript, through the use of pharmacological modulation of cytosolic Ca^2+^, we determine the properties of the calcium signal that enhances autophagy and the likely step in the induction pathway that is being affected.

## Materials and methods

### Cell culture

All cell culture solutions were obtained from Invitrogen and all plastic-ware was obtained from Nunc, unless otherwise stated. Human embryonic kidney (HEK) 293 cells were maintained in Dulbecco’s Modified Eagle Medium (DMEM) supplemented with 10% fetal bovine serum (FBS), 1% penicillin (10,000 IU) and 1% streptomycin (10,000 μg/ml). HEK 293 stably transfected with GFP-DFCP1 [8] or GFP-DFCP1 + RFP-ATG9 were cultured in media identical in composition to wild-type media, except for the addition of 400 μg/ml Geneticin (G418, Gibco, 11811031) or 400 μg/ml G418 and 100 μg/ml Zeocin (Invivogen, ant-zn-05), respectively. WT and KO ATG9 and ULK1/2 MEFs were kindly provided by Sharon Tooze. WT and KO ATG13 and FIP200 were kindly provided by Noboru Mizushima and Jun-Lin Guan, respectively. We have validated the absence of the relevant proteins in those MEFs in a previous publication from our lab [7]. MEFs were cultured in the same medium as HEK 293 cells but supplemented with 1x non-essential amino acids (Gibco, 11140050). All cells were kept at 37°C in a 5% CO_2_ humidified atmosphere.

### Antibodies and reagents (catalogue numbers are in parenthesis)

ATP, bafilomycin A1 (B1794), BAPTA-2 AM (16609), CPA (C1530), Fura-2 AM (F0888), PP242 (P0037), STO-609 (S1318), thapsigargin (T9033) and wortmannin (W1628) were purchased from Sigma. A23187 (BML-CA101-0001) was purchased from Enzo. EGTA-AM (E1219) was purchased from Invitrogen. The following primary antibodies were used for immunofluorescence following either formaldehyde fixation [anti-ATG13 (Millipore, MABC46); anti-ATG16L (MBLI, PM040); anti-ATG101 (Sigma, SAB200175); anti-FIP200 (ProteinTech Europe, 17250-1-IP); anti-LAMP1 (Pharminingen, 34021A); anti-phospho-ATG13 Ser318 (Rockland, 600-401-C49), anti-ULK1 (Sigma, A7481); anti-WIPI2 (Bio-Rad, MCA5780GA)] or methanol fixation [anti-LC3B (Sigma; L7543)]. The following primary antibodies were used for immunoblotting: anti-GAPDH (Biogenesis, 4699-9555); anti-ULK1 (Santa Cruz, SC33182); anti-phospho-ULK1 Ser757 (Cell Signalling Technology; 6888S); anti-LC3B (Novus Biologicals, NB 100-2331); anti-β-COP (hybridoma clone M3A5; a kind gift from the late Dr Thomas Kreis). The following secondary antibodies were used for either immunoblotting or immunostaining and were purchased from Jackson Immuno Research: goat anti-rabbit/mouse IgG HRP conjugate (115-035-146; 111-035-144), goat anti-rabbit/mouse IgG Cy2-(115-225-146; 111-225-144) or RITC-(115-025-062; 111-025-144) conjugated secondary antibodies.

### Cell culture treatments

Cells were amino acid-starved in 3 ml of starvation medium (140 mM NaCl, 1 mM CaCl_2_, 1 mM MgCl_2_, 5 mM glucose and 20 mM HEPES, pH 7.4) for 1 h at 37°C in a 5% CO_2_ humidified atmosphere.

Live-cell imaging was performed in Ham’s F-12 nutrient mix (Thermo Scientific 11765054) supplemented with 15 mM HEPES, 1 mM CaCl_2_, 10% FBS (F12-medium).

### Cell transfections

For siRNA knockdown, we used predesigned oligonucleotides from Horizon Discovery (SMARTpool) to reduce expression levels of FIP200. Cells were transfected using DharmaFECT 1 (Horizon Discovery) and examined 72 h later. HEK 293 cells stably-expressing GFP-DFCP1 and RFP-ATG9 were generated by transfection of HEK 293 GFP-DFCP1 cell with RFP-ATG9 using X-tremeGENE 9 (Roche) as per the manufacturer’s instructions. Transfected clones were then selected for by growth in medium containing 800 μg/ml G418 and 100 μg/ml Zeocin for 12 days.

### Immunofluorescence microscopy

Following the appropriate treatments, cells grown on glass coverslips in 12-well plates were fixed by washing once in 3.7% formaldehyde and then incubating in 3.7% formaldehyde for 20 min at room temperature. The cells were then washed twice in DMEM/HEPES and incubated for 20 min in the same solution to quench excess formaldehyde. Alternatively, fixation was performed by treatment with 100% ice-cold methanol (stored at −20°C) for 10 min on ice. The cells were then blocked for a minimum of 1 h in 2% (w/v) BSA/PBS on a shaker before staining with the appropriate antibodies. Immunostaining of formaldehyde-fixed cells required initial permeabilization of samples using NETgel [150 mM NaCl, 5 mM EDTA, 50 mM Tris-Cl pH 7.4, 0.05% Nonidet P-40 (NP40, Roche, 11754599001), 0.25% gelatin and 0.02% Na azide] supplemented with 0.25% NP-40 for 10 min on a shaker at room temperature. Following permeabilization, samples were washed twice in NETgel to remove excess detergent before staining with the appropriate antibody. NETgel was then used for all antibody dilutions and washes. Immunostaining of methanol fixed cells was performed in a similar manner to that described above but did not require the permeabilization step because methanol fixed and permeabilized the cells at the same time. Thereafter, 2% BSA (w/v)/PBS was used for all washes and antibody dilutions.

### Preparation of cell lysates for western blotting

Following appropriate treatments, cells growing on tissue culture plates were placed on ice and washed 3 times in ice cold PBS. Cells were then lysed in 100 μl of lysis buffer (50 mM Tris-Cl pH 8.0, 50 mM KCl, 10 mM EDTA, 0.6 mM PMSF, 1 μg/ml trypsin inhibitor 1% NP-40) for 16 min, then collected using a rubber cell scraper. After centrifugation at 12,000 x g at 4°C for 10 min, the cell debris was discarded and the supernatant transferred to a fresh 1.5 ml microcentrifuge tube and stored at −20°C. Western blotting was performed using standard methods. Samples were processed for SDS-PAGE and analysed by standard western blotting techniques.

### Loading of cells with Fura-2 AM

Stock solutions of Fura-2 AM (1 mM) were prepared in DMSO (Sigma) containing 20% F-127 Pluronic (Invitrogen). Stock solutions were maintained at -20°C in the dark. For loading, cells were incubated with 1 μM Fura-2 AM diluted in F-12 medium for 30 min at 20°C. Intracellular Fura-2 AM was then allowed to de-esterify by replacement of the imaging buffer with fresh buffer for 30 minutes. The Fura-2 AM ratio (340/380) used as a measure of [Ca^2+^]_i_ changes was obtained from the background-subtracted emission fluorescence intensity of the dye when excited at 340 nm and 380nm.

### Epifluorescence microscopy

Images of fixed cells were acquired using a Zeiss Axio Imager D2 wide-field epifluorescence microscope, using a 63x 1.4 NA objective, AxioCam HR CCD camera (Zeiss) and HXP 120C metal halide light source (Zeiss).

### Live-cell imaging

#### Fura-2 AM

Cells were imaged on an Olympus IX81 wide-field microscope equipped with an Olympus 40x/340 1.35 NA oil immersion lens and fitted with a full enclosure Solent Scientific environment chamber and maintained at 37°C during imaging. An Olympus MT-20 xenon/mercury illumination unit was used to provide excitation light and a Hamamatsu Orca-ER CCD camera captured image data. Fura-2 AM was imaged using alternate excitation filters at 340/26 (Semrock) and 387/11 (Semrock) and a 525/40 emission filter (Chroma Technology). The system was controlled using Olympus CellR software.

#### GFP-DFCP/RFP-ATG9

Cells were imaged using a system configured around a Nikon Ti-E microscope equipped with a 60x 1.4 N.A. objective (Nikon) and controlled using NIS Elements software (VersionXX). Excitation light was provided by a SpecraX LED illuminator (Lumencor) and emitted light was collected using a Hamamatsu Flash 4.0 sCMOS camera. The light path included 410/504/582/669-Di01 and Di01-R442/514/561 dichroic mirrors (Semrock) and excitation emission filter pairs for GFP (480/10 (ex) 525/30 (em)) and mRFP (560/25 (ex) 607/36 (em) all from Semrock), which were controlled using a filter wheel (Sutter Instruments).

### Super-resolution microscopy

Super-resolution images were acquired using a Nikon N-SIM structured illumination microscope D-STORM comprising a Nikon Ti-E microscope, Nikon 1.49 NA 100x oil immersion objective, Andor iXon 897 EM-CCD camera, MBP Communications 643 nm laser (300 mW), Coherent Sapphire 561 nm laser (150 mW), Coherent Cube 405 nm laser (100 mW) and Semrock LF405/488/561/635-A-000 multiband filter set. Labelled samples were mounted into custom imaging rings and immersed in 500 μl STORM imaging buffer comprising (all from Sigma) catalase (1 μg/ml), TCEP (0.8 μM, glucose (40 mg/ml), glucose oxidase (50 μg/ml), glycerine (12.5%), KCl (1.25 mM), Tris-HCl (1 mM) and MEA-HCl (100 nM). The imaging system was controlled using Nikon Elements software. Images 256×256 pixels (0.16 μm/px) were captured at a frame rate of 55 frames per second with the camera conversion gain set at 2.4 and the EM gain typically set at 100. At least 15,000 frames were captured per field of view, with the laser illumination switching alternately for dual labelling experiments. Nikon Elements (NIS Elements Version 4.13) software was used for image acquisition and super resolution image reconstruction

### Quantitation of puncta

Quantification and determination of colocalization of puncta was performed using the Spots Detection function of Imaris software (Bitplane/Andor). The measurements were made on 10 randomly selected fields of view.

Live-cell imaging videos were analysed with Fiji Open Source software. We define as an independent event all the frames that correspond to the formation and collapse of one GFP–DFCP1 containing particle, starting and finishing with the frames in which the fluorescence of the GFP–DFCP1 particle is emerging clearly above the fluorescence of the cytosolic GFP–DFCP1. Frames corresponding to time points before the beginning, or after the end of a particular event were carefully scanned, and events corresponding to particles moving in and out of focus were excluded from the analysis.

### Statistical analysis

Statistical analysis was performed on data where indicated in the figure legends. All statistical analysis was performed using GraphPad Prism software. Data obtained from at least two independent experiments were used in all statistical tests. Before running ANOVAs and t-tests, assumptions for parametric tests were checked. All multiple comparison tests were carried out using a Sidak post hoc test. The type of statistical test used can be found in the figure legend. Statistical significance was determined as follows: * p<0.05; **p<0.01; ***p<0.001;****p<0.0001. All data are presented with error bars that represent mean ±SEM. Data obtained with the help of Dr. Anne Segonds-Pichon (Babraham Institute).

## Results

### Ca^2+^ release from intracellular stores induced by A23817 and thapsigargin cause translocation of early autophagic proteins to pre-autophagosomal puncta

To identify novel compounds that induce omegasome formation, we performed a chemical screen with the <u>L</u>ibrary <u>O</u>f <u>P</u>harmacologically <u>A</u>ctive <u>C</u>ompounds (LOPAC^1280 TM^) and using HEK 293 cells stably expressing GFP-DFCP1 as our readout. We looked for chemicals that induced DFCP1 puncta in normal medium. From this screen, we identified A23187, a mobile ion-carrier (an ionophore) that induces an intracellular Ca^2+^ elevation by conveying Ca^2+^ from the extracellular medium and from ER Ca^2+^ stores into the cytosol. A23187 stimulated similar omegasome induction to that seen with thapsigargin (Tg, an agent that induces cytosolic Ca^2+^ elevation), which we had previously identified as a potent activator of omegasome responses (Fig 1A and B). In parental HEK 293 cells not expressing GFP-DFCP1, both of these compounds induced the formation of endogenous WIPI2-positive puncta (WIPI2 is an important effector of PI3P during the autophagy induction step [44]) to levels similar to those seen during canonical autophagy induced either by starvation or by chemical inactivation of mTOR (Fig 1 C-E). Interestingly, puncta formed by ATG16L (a later autophagy protein interacting with WIPI2 [49]) were less in number, and only partially co-localised with WIPI2 (Fig 1C-E). This suggested that the response elicited by these compounds primarily affects the early autophagic intermediates and is insufficient to induce a complete autophagic response. This was further confirmed by analysis of LC3 lipidation assays using immunoblots: whereas the lipidation of LC3 upon mTOR inhibition (treatment with PP242) was strong and further enhanced by bafilomycin A1, it was very close to background levels for the two calcium-related treatments (Fig 1F, note levels of LC3-II which is the lipidated form). This early calcium-related autophagic response was sensitive to PI3P inhibition by wortmannin (Fig 1G for endogenous WIPI2 and H for GFP-tagged WIPI2), showing that it is acting through the canonical PI3P-dependent autophagic pathway.

**Figure 1.**
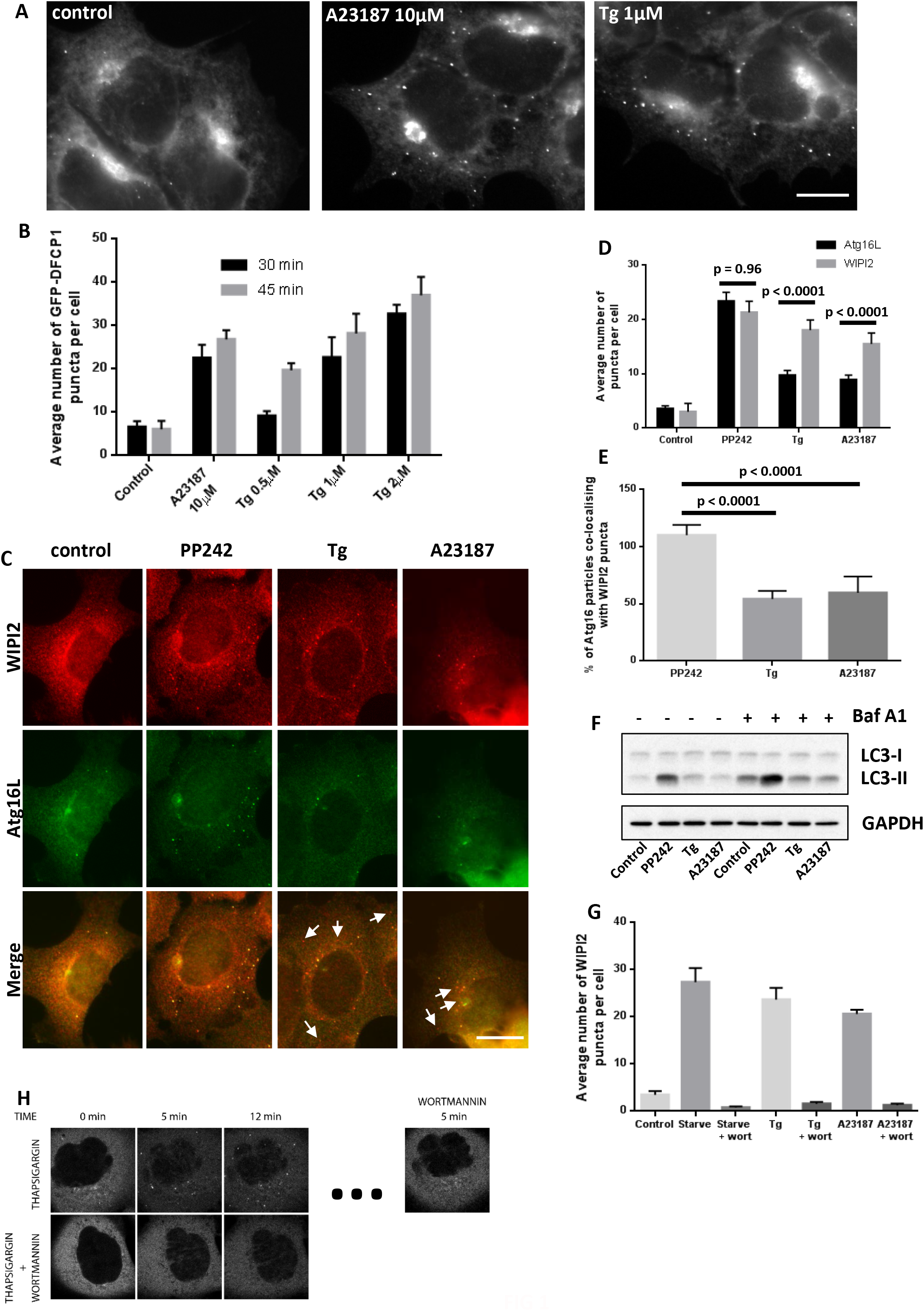
Elevation in cytosolic [Ca^2+^] causes formation of omegasome-related intermediates but does not result in a full autophagy response. (A) HEK 293 cells stably expressing GFP-DFCP1 were treated with A23187 or Tg for 30 min and then fixed and examined by fluorescence microscopy. (B) Cells, as in A, were treated with 10 µM A23187 or a range of concentrations of Tg, as indicated, in fed conditions for 30 and 45 min (n = 2, ± SEM). The number of GFP-DFCP1 puncta per cell was quantitated and plotted. (C-E) HEK 293 cells were treated for 30 min with PP242 (1 μM), Tg (1 μM) or A23187 (10 μM), fixed and co-immunostained for endogenous WIPI2 and ATG16L. Tg- and A23187-induced WIPI2 puncta often did not co-localise with ATG16L (white arrows in C). Quantitation of the individual WIPI2 and ATG16L puncta (n = 3, ± SEM) is shown in D whereas the percentage of ATG16L-positive puncta that co-localise with WIPI2-positive puncta is shown in E (n = 3, ± SEM). (F) Immunoblot showing LC3-II levels after 30 min of treatment with PP242 (1 μM), Tg (1 μM) or A23187 (10 μM) +/- bafilomycin A1 (100 nM). (G) HEK 293 cells were starved (1 h treatment), or treated with 1 µM Tg or 10 µM A23187 (both 30 min treatment) in the presence or absence of 66 nM wortmannin as indicated. Cells were fixed and immunostained for endogenous WIPI2 and the average number of WIPI2 puncta per cell was quantitated and plotted (n = 2, ± SEM). (H) HEK 293 cells expressing GFP-WIPI2b were imaged live in the presence of 200 nM Tg and then 200 nM wortmannin was added as indicated (top panels, selected frames from the movie sequence are shown). Alternatively, wortmannin and Tg were added together and the cells imaged for 8.5 min (bottom panel).

### Sustained, elevated basal cytosolic calcium levels are required for omegasome formation

Although the SERCA pump inhibitor Tg acts primarily on the ER and the ionophore A23187 on the plasma membrane, both agents ultimately lead to Ca2+ flux across both the ER and plasma membrane resulting in an elevation in cytosolic Ca2. To address how these compounds modulate autophagic induction, we performed live-cell imaging experiments to simultaneously observe omegasome formation (using GFP-DFCP1 as a reporter) and cytosolic calcium levels (loading with Fura-2 acetoxymethyl ester [Fura-2 am]) in the same cells (Fig 2). Tg and A23187 both elicited an increase in omegasome formation that always began subsequent to the initial cytosolic Ca^2+^ elevation and after Ca^2+^ levels had reached a plateau above that of the initial baseline prior to intervention (Fig 2B, C and E). Notably, cell stimulation with ATP, which induced a robust transient Ca^2+^ increase that rapidly returned to the initial basal state, did not produce such an omegasome response (Fig 2D). Further, no omegasome response was ever detected even when increased concentrations of ATP were applied once or were repetitively applied to increase the amplitude or prolong the elevation in cytosolic Ca^2+^ levels respectively (Fig S1). In addition, use of the reversible SERCA pump inhibitor cyclopiazonic acid (CPA), which generates a Ca^2+^ increase similar to that induced by Tg but the action of which can be rapidly reversed, returning Ca^2+^ levels to baseline, showed that a sustained elevation of Ca^2+^ levels is required for an omegasome response (Fig S2). Taken together, these data suggest that irrespective of the magnitude of the initial cytosolic Ca^2+^ peak induced by cell stimulation, a small but sustained elevation of basal cytosolic Ca^2+^ levels is necessary for triggering the omegasome response. The cumulative data from these treatments showing the extent of the Ca^2+^ elevation are plotted in Fig 2E.

**Figure 2.**
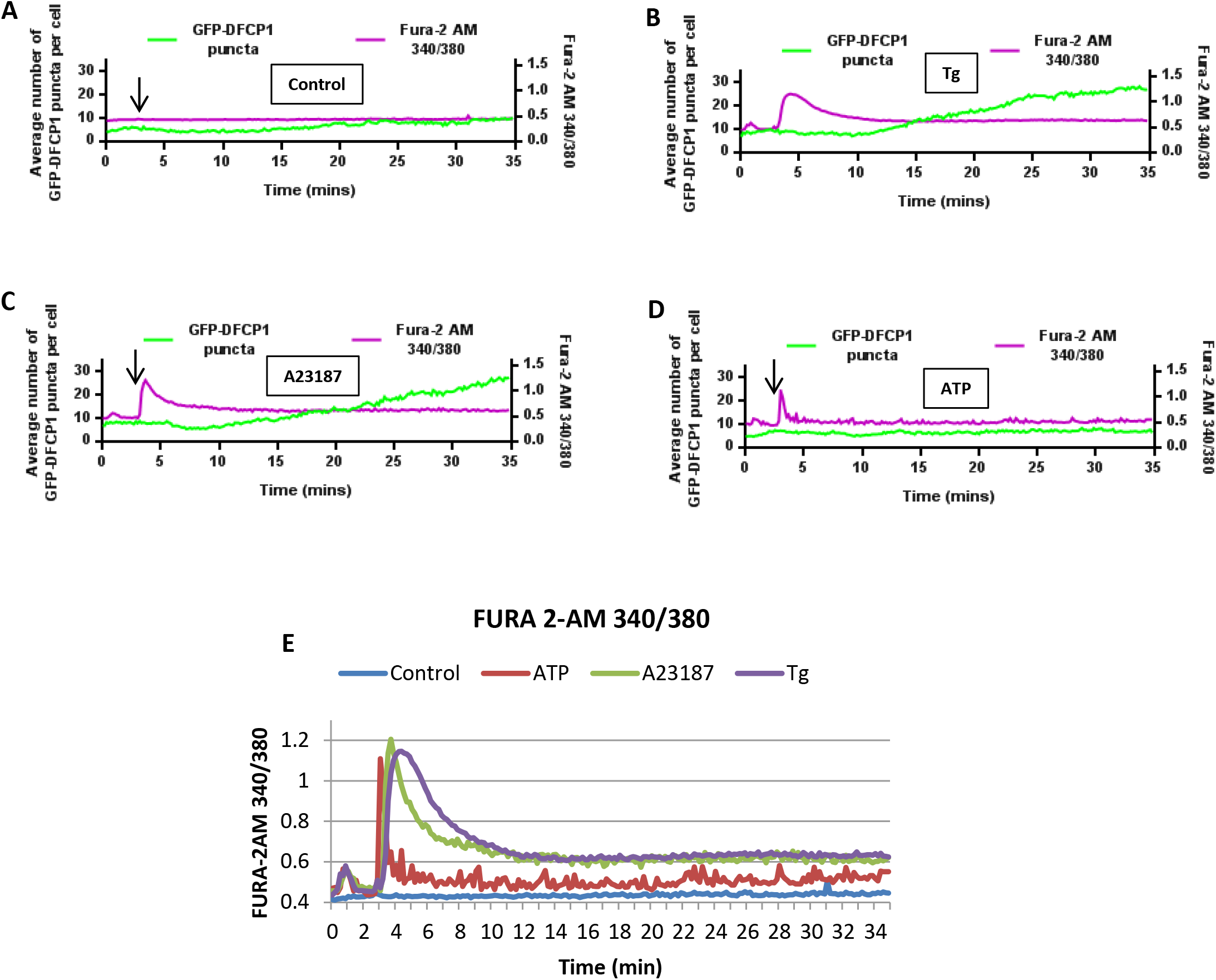
Dynamics of omegasome formation in response to elevation of cytosolic [Ca^2+^]. HEK 293 cells stably expressing GFP-DFCP1 were loaded with Fura-2 AM and imaged live following 4 treatments: (A) DMSO, (B) 1 μM Tg, (C) 10 μM A23187, (D) 2.5 μM ATP. All additions occurred 3 min after the start of imaging (arrows). Individual cells were then identified and the GFP-DFCP1 puncta number and Fura-2 AM 340/380 ratio measured in the same cell. Each graph represents the average values for 30 cells collected over 4 experiments in 2 separate imaging sessions. (E) Overlay of the Fura-2 AM ratio for graphs A-D.

Although we have restricted our Ca^2+^ elevating treatments to relatively short times, our data are in line with those from Hoyer-Hansen et al. who identified prolonged Ca^2+^ elevation (24h) as an inhibitor of mTOR and an activator of Ca^2+^/calmodulin-dependent kinase kinase-β and AMP-activated protein kinase, which together induce autophagy [17]. Our results are independent of this pathway, however, because treatment with the CaMKK inhibitor STO-609 had no impact on omegasome formation caused by Tg or A23187, showing that this shorter-term Ca^2+^response is AMPK-independent (Fig S3). Our data that the Ca^2+^-dependent response is independent of AMPK are consistent with those of Grotemeier et al. [25] who showed that Ca^2+^ elevation induces WIPI1 puncta in the absence of AMPK.

### Elevation in cytosolic Ca^2+^ causes translocation of the entire ULK complex to pre-autophagosomal puncta and phosphorylation of ATG13 on Ser318

During the autophagic response, the ULK complex (composed of ULK1, FIP200, ATG13 and ATG101) translocates to punctate structures that associate with the ER and with ATG9 vesicles to initiate autophagosome formation via PI3P-enriched omegasomes [50]. As this translocation lies upstream of the PI3P-dependent step that we originally observed to be sensitive to Ca^2+^ elevation, we investigated whether cytosolic Ca^2+^ changes are also acting at this early initiation step. All four components of the ULK complex formed foci upon pharmacological increases in cytosolic Ca^2+^ to a similar level to that produced by the mTOR inhibitor, PP242 (Fig 3A and B). Whereas similar in number, these foci appeared smaller than those formed by PP242 (see also below). Interestingly, phosphorylation of ATG13 on Ser318, the site that is regulated by inactivation (de-phosphorylation) of ULK1 downstream of mTOR inactivation [51], was elevated by increased cytosolic Ca^2+^ levels but without an accompanying de-phosphorylation of ULK1 (Fig 3C-G).

**Figure 3.**
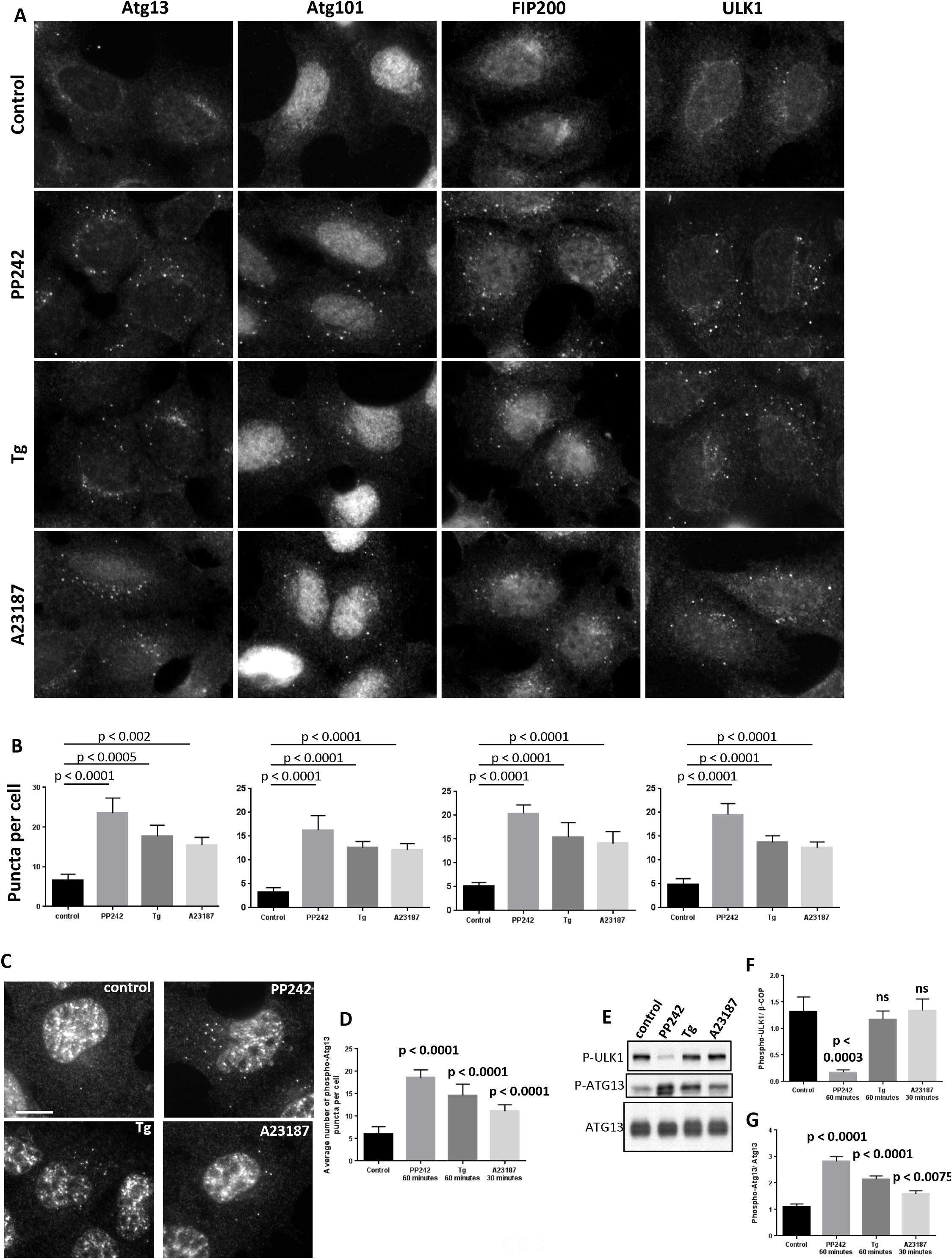
Elevation of cytosolic [Ca^2+^] induces the translocation of the entire ULK complex to pre-autophagosomal structures and the phosphorylation of ATG13. (A) HEK 293 cells were treated with 1 μM PP242, 1 μM Tg or 10 μM A23187 for 30 min, fixed and immunostained for endogenous ATG13, ATG101, FIP200 or ULK1. (B) Quantification of average puncta per cell of samples from A (n = 3, ± SEM). (C) HEK 293 cells were treated with 1 μM PP242 (1h), 1 μM Tg (1 h) or 10 μM A23187 (45 min), fixed and immunostained for endogenous phospho-Atg13 Ser318. (D) Quantitation of data in C (n = 3, ± SEM). (E) Cells treated as in C were lysed and the lysates immunoblotted for phospho-ULK1 Ser757, phospho-Atg13 Ser318 and ATG13. (F, G) Levels of phospho-ULK1 Ser757 normalised to β cop (not shown) and phospho-Atg13 Ser318 normalised to non phosphorylated ATG13 as shown in E are plotted (n = 3, ± SEM).

### Suppression of cytosolic Ca^2+^ inhibits translocation of the entire ULK complex to pre-autophagosomal puncta induced by starvation, and reduces phosphorylation of ATG13 on Ser318

Given the significant response of the ULK complex to Ca^2+^ elevation, we next examined whether cytosolic Ca^2+^ chelation would block autophagic induction at the level of the ULK complex as well (Fig 4). Induction of autophagy by PP242, as measured by puncta formation of all of the ULK complex members, was supressed by treatment with the cell permeable Ca^2+^ chelator BAPTA-2 AM, (Fig 4 A, B). Similarly, the increase in phosphorylation of ATG13 at Ser318 caused by autophagy induction was completely suppressed by treatment with BAPTA-2 AM (4C-G). BAPTA-2 AM treatment also blocked later steps in autophagy as measured by LC3 lipidation and LC3 puncta formation (Fig S4). Because BAPTA-2 AM can chelate zinc as well as calcium, we used the zinc chelator TPEN to examine if it would also affect the autophagic response. Although TPEN inhibited the DFCP1 response (DFCP1 contains two FYVE domains that require zinc for functionality), it did not inhibit the response of any of the other proteins such as members of the ULK complex or WIPI2 (data not shown).

**Figure 4.**
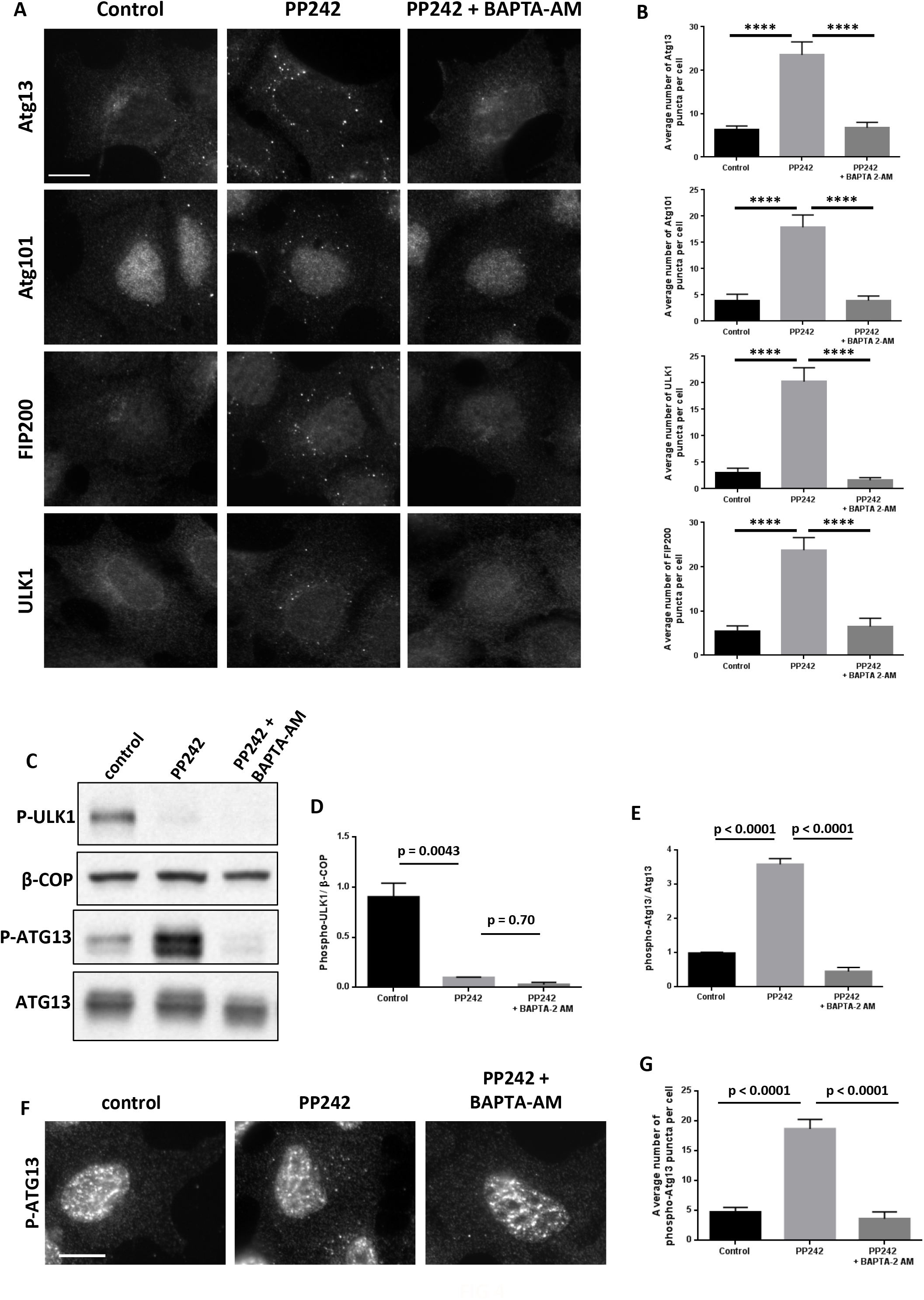
Chelation of cytosolic Ca^2+^ with BAPTA-2 AM inhibits starvation-induced translocation of the ULK complex to pre-autophagosomal structures and the phosphorylation of ATG13. (A) HEK 293 cells were treated with 1 μM PP242 alone or in the presence of 10 μM BAPTA-2 AM (BAPTA-2 AM was added 5 min before PP242). Cells were fixed and immunostained for endogenous ATG13, ATG101, FIP200 or ULK1. Scale bar is 10 μm. (B) Levels of average puncta per cell from A (n = 3, ± SEM). (C) HEK 293 cells were treated as in A, lysed and immunoblotted for phospho-ULK1 Ser757, and phospho-ATG13 Ser318. (D, E) Levels of phospho-ULK1 Ser757 and phospho-ATG13 Ser318 against the indicated controls are plotted (n=3 ± SEM). (F) Cells treated as in A were immunostained for phospho-ATG13 Ser318. Scale bar is 10 μm. (G) Levels of average phospho-ATG13 Ser318 puncta per cell from F (n = 3, ± SEM).

Intracellular calcium can also be chelated with cell permeable EGTA-AM albeit with slower kinetics than BAPTA-2 AM. As such, while EGTA prevents global Ca2+ elevations, it does not affect local Ca2+ elevations generated, at for example, the mouth of a Ca^2+^ channel [52]. This compound did not block omegasome formation (Fig 5A) or ULK complex translocation (not shown). In addition, unlike BAPTA-2 AM, EGTA-AM did not alter phosphorylation state of ATG13 during autophagy induction (Fig 5B, C). These findings suggest that a fast and localised Ca^2+^ signal is responsible for these effects [53].

**Figure 5.**
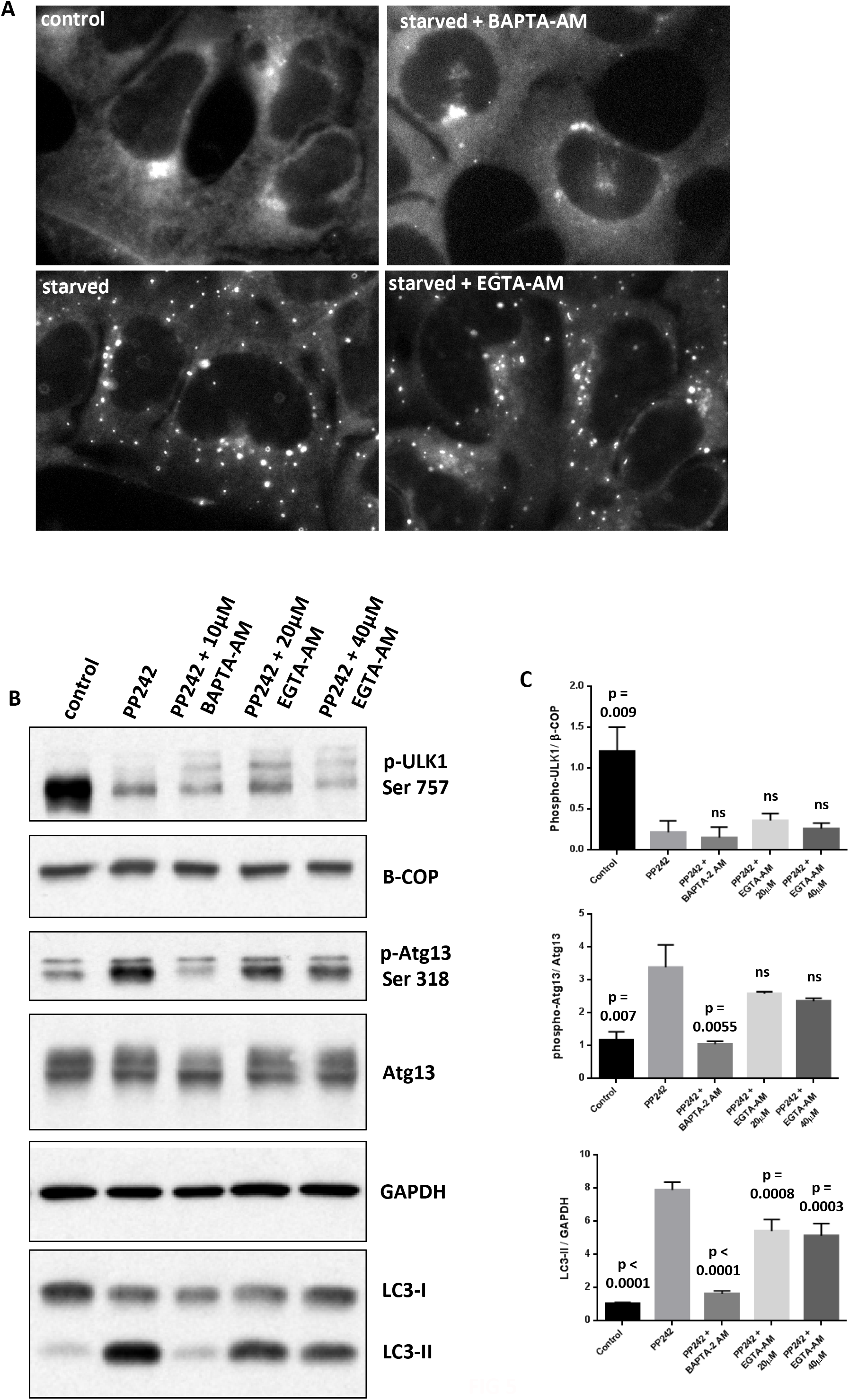
Chelation of cytosolic Ca^2+^ with EGTA-AM does not affect starvation-induced responses. (A) HEK 293 cells stably expressing GFP-DFCP1 were starved in the presence or absence of 10 µM BAPTA-2 AM and or 40 µM EGTA-AM, and were examined by fluorescence microscopy. Scale bar is 10 μm. (B) HEK 293 cells were treated with PP242 in the presence or absence of BAPTA-2 AM or EGTA-AM as indicated, lysed and immunoblotted for phospho-ULK1 Ser757, phospho-Atg13 Ser318, Atg13 and LC3. BAPTA-2 AM and EGTA-AM were added 5 min before PP242. (C) Levels of the indicated proteins from B are shown (n = 3, ± SEM).

Since Tg and A23187 promote both Ca2+ release from intracellular stores and Ca2+ entry across the plasma membrane, our data do not establish which is the source of Ca^2+^ (either intracellular or extracellular) and thus which is required for eliciting an early autophagy response. We therefore wanted to examine whether chelation of extracellular Ca^2+^ and thus prevention of Ca2+ entry into the cell would be inhibitory to autophagy. When cells were starved in the absence of any extracellular Ca^2+^, the expected autophagic response was induced (data not shown). This finding is consistent with the notion that an intracellular Ca^2+^ source provides the physiological Ca^2+^ signal for autophagy induction.

### Ca^2+^-induced early pre-autophagosomal puncta are functionally and morphologically related to autophagy intermediates

A characteristic of early autophagy intermediates, such as omegasomes, is that they almost always emerge in association with a subset of ATG9-positive vesicles [54]. We therefore examined if the autophagic structures that are induced through increases in cytosolic Ca^2+^ depend on the presence of ATG9. In MEF cells devoid of ATG9, Ca^2+^-induced WIPI2 puncta were severely reduced (Fig 6A and B). In HEK 293 cells expressing the omegasome component GFP-DFCP1 and RFP-ATG9, the large majority of newly formed Ca^2+^-induced omegasomes were in association with ATG9 vesicles, identical to the puncta forming during induction of autophagy by mTOR inhibition using PP242 (Fig 6C). Furthermore, the lifespan of GFP-DFCP1 puncta formed by treatment with the mTOR inhibitor PP242 was comparable to that formed by the two compounds that elevate cytosolic Ca^2+^ (Fig 6D and E). To examine the morphology of these Ca^2+^-induced puncta containing FIP200 and ATG13 at higher resolution, we used super resolution Stochastic Optical Reconstruction Microscopy (STORM). We found that, in comparison to the structures formed during mTOR inhibition, the Ca^2+^-induced puncta were more compact and less elaborate (Fig 6F).

**Figure 6.**
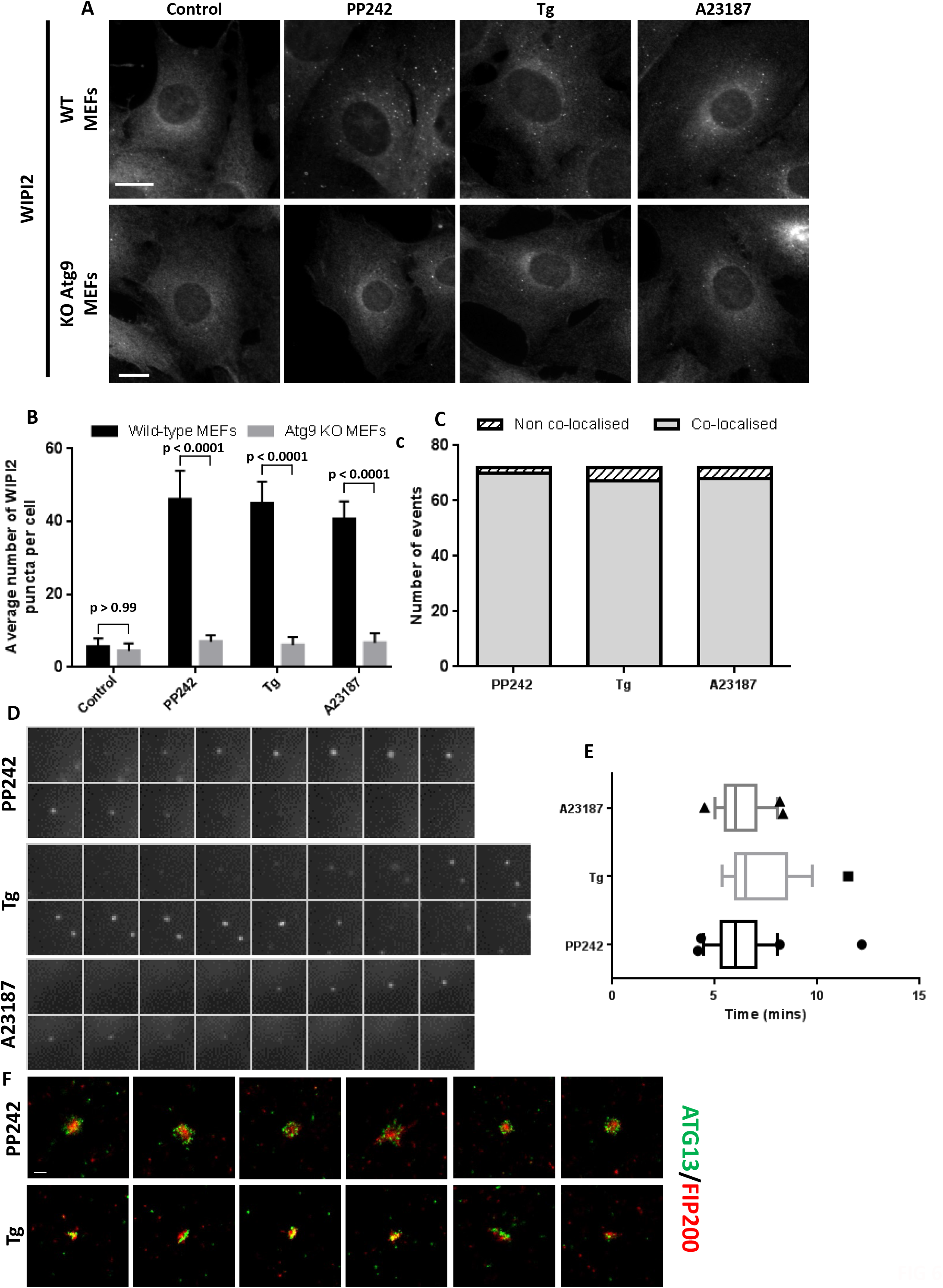
Ca^2+^-induced pre-autophagosomal structures require ATG9, have similar life-spans to canonical starvation-induced structures but are smaller and less elaborate. (A) WT MEFs or Atg9 KO MEFs were treated for 30 min with 1 μM PP242, 1 μM Tg or 20 μM A23187, fixed and immunostained for endogenous WIPI2. (B) Average number of WIPI2 puncta per cell from A (n = 3, ± SEM). Scale bar is 10 μm. (C) Live-cell imaging of HEK 293 cells stably expressing GFP-DFCP1 and RFP-ATG9 and treated with: 1 μM PP242, 1 μM Tg or 10 μM A23187. The number of newly formed GFP-DFCP1 puncta that co-localised with an RFP-ATG9 vesicle present at the first frame of each event is plotted. These measurements are derived from n = 72 cells. (D) Montages representing typical life span of a GFP-DFCP1 vesicle formed after treatment with 1 μM PP242, 1 μM Tg or 10 μM A23187. (E) Box plot representing the average life span of GFP-DFCP1 puncta shown in D. Data derived from PP242 (n = 26 puncta), Tg (n = 17 puncta) or A23187 (n = 23 puncta) from 2-4 experiments. Error bars represent 10-90 percentile. (F) STORM super-resolution microscopy of FIP200 and ATG13 puncta induced by PP242 or Tg. HEK 293 cells were treated with 1 μM PP242 or Tg for 60 or 30 min, respectively. Cells were then fixed, immunostained and imaged using STORM. Scale bar = 0.2 µm.

### Formation of Ca^2+^-induced early pre-autophagosomal puncta does not require ULK1/2

As previously reported, MEF cells that lack both ULK1 and ULK2 are unable to activate autophagy in response to starvation or chemical inhibition of mTOR [55]. Interestingly, the autophagic structures induced by Ca^2+^ elevation did not require the presence of ULK1 and ULK2 although those proteins were still required as expected for an autophagic response to the mTOR inhibitor PP242 (Fig 7A and B). In contrast, the loss of ATG13, either in MEFs or in HEK 293 cells, completely inhibited the Ca^2+^ - induced response (Fig 7 C, D). These ATG13-negative cells also showed a complete inhibition of autophagy induced by the mTOR inhibitor PP242 (Fig 7 C, D) and by amino acid starvation (not shown). Similarly, the loss of FIP200 in MEFs fully abrogated the formation of autophagic structures induced by mTOR inhibition or Ca^2+^elevation (shown for Tg in Fig 7 E, F). These results suggest that the Ca^2+^-induced formation of early pre-autophagosomal structures differs mechanistically from an early autophagy response in that it is independent of ULK1/2 but dependent on FIP200, ATG13 and ATG9.

**Figure 7.**
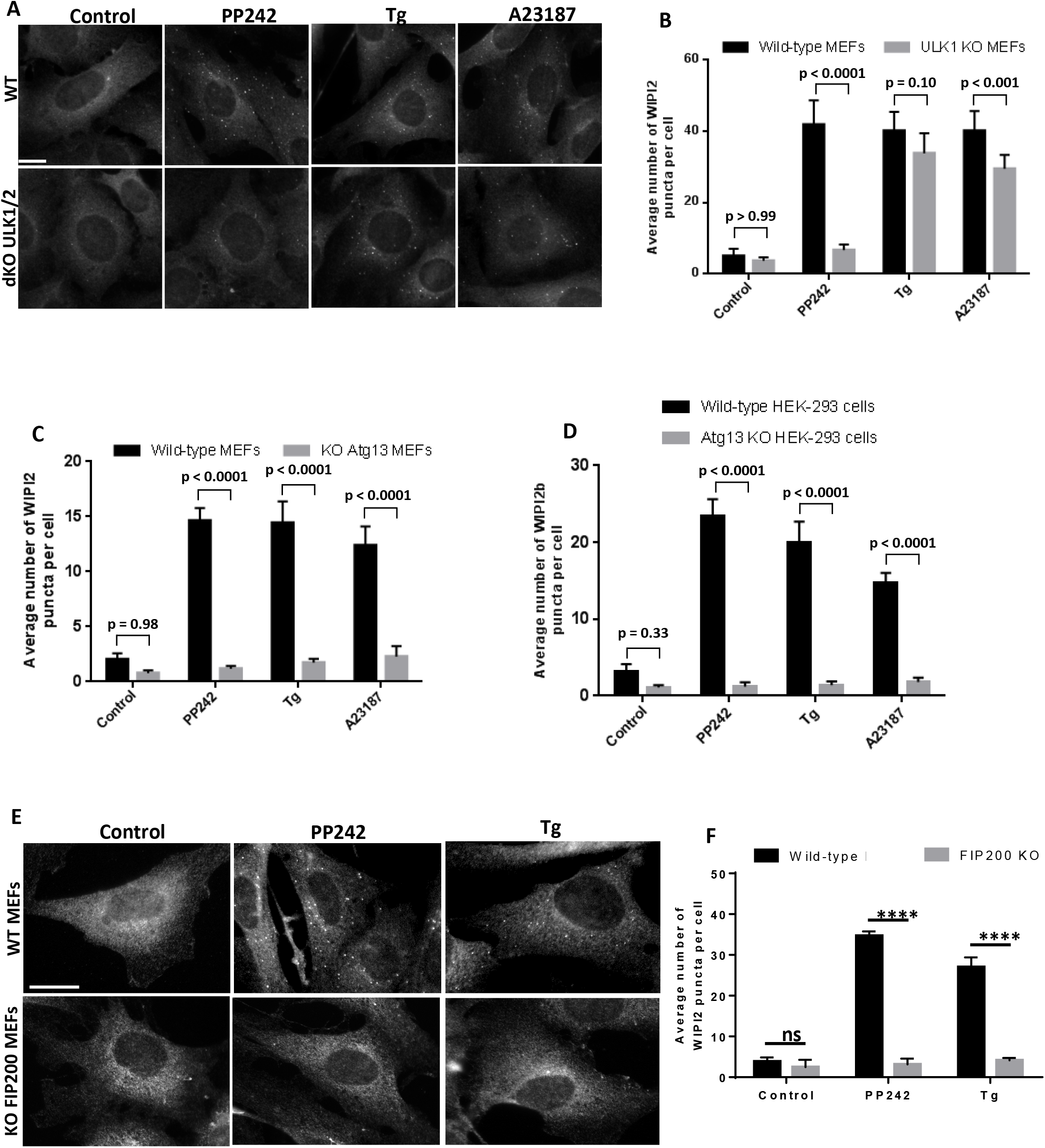
The Ca^2+^-induced pre-autophagosomal structures can form in the absence of ULK1 and ULK2 but require ATG13, and FIP200. (A) Wild-type MEFs or those with a double knockout for ULK1 and ULK2 were treated for 30 min with 1 μM PP242, 1 μM Tg or 20 μM A23187, fixed and immunostained for endogenous WIPI2. (B) The average number of WIPI2 puncta per cell is shown (n = 3, ± SEM). (C) Wild-type MEFs or those MEFS from ATG13 knockout mice were treated as in A, and the average number of WIPI2 puncta per cell is plotted (n = 3, ± SEM). (D) Wild-type HEK 293 cells or those carrying a CRISPR-mediated deletion of ATG13 were treated as in A and the average number of WIPI2 puncta per cell is plotted (n = 3, ± SEM). (E) Wild-type MEFs or those with a knockout for FIP200 were treated as in A. (F) The average number of WIPI2 puncta per cell for the samples in E is shown (n = 3, ± SEM). Scale bar is 10 μm.

Given the strong requirement of the Ca^2+^-induced puncta on FIP200 and ATG13, we then investigated if each of these proteins on their own could respond to Ca^2+^ modulation (Fig S5). Knockdown of FIP200 protein using siRNA prevented the formation of ATG13-puncta after Tg treatment. Similarly, elimination of the ATG13 protein using CRISPR also prevented the FIP200 translocation to puncta induced by Ca^2+^. From these data, we conclude that ATG13 and FIP200 respond as a single unit to a cytosolic Ca^2+^ increase during autophagy induction.

## Discussion

In this work, we have explored the potential mechanisms by which Ca^2+^ elevation stimulates early steps during autophagy induction and Ca^2+^ chelation prevents those steps. By simultaneously imaging omegasome dynamics and Ca^2+^ levels after various treatments, we have shown that a rather moderate elevation in basal cytosolic Ca^2+^ levels within 15 min is sufficient to stimulate autophagy, with the provenance of the Ca^2+^ (coming either from internal stores or from the extracellular medium) not being important. This Ca^2+^ -dependent omegasome response is not synchronous with the initial Ca^2+^elevation but rather with the time when the Ca^2+^signal has dropped from its peak and has stabilised at a new, higher than the initial, basal level. It is known that the dynamic range of intracellular Ca^2+^ signals is large, ranging from 100 nM in the resting state to tens of micromolar near the Ca^2+^ source after stimulation [56]. However, the majority of such signals are oscillatory with a periodicity of seconds and only in selected cellular conditions they can stay elevated for longer times. In this sense, the fact that early autophagy proteins become mobilised only after a sustained Ca^2+^ elevation would ensure that the autophagy pathway would not be activated by any momentary Ca^2+^ fluctuation underlying normal Ca^2+^ signalling.

The lag time between initial Ca^2+^ elevation and autophagic response suggests that an activation or assembly of some machinery must take place once the new elevated plateau of Ca^2+^ levels is reached. To identify what this machinery maybe we examined the earliest autophagy-specific complex consisting of the kinase ULK1 (or ULK2) and the associated proteins FIP200, ATG13 and ATG101. All four translocated to Ca^2+^-induced puncta, and all were inhibited from translocation when Ca^2+^ was depleted during an autophagic stimulus. We also discovered that the phosphorylation of ATG13 at serine 318, known to be dependent on ULK1 [51], completely tracked both the stimulatory and the inhibitory Ca^2+^ effect as outlined above. Based on these observations, a straightforward mechanism would be that Ca^2+^ regulates the kinase activity of ULK1. However, this cannot be the case since we also discovered that ULK1 and ULK2 are completely dispensable for the Ca^2+^-dependent response. To eliminate the possibility that an alternative kinase was phosphorylating ATG13 in the ULK1/ULK2 KO MEFs, we examined levels of phospho-ATG13 Ser318 following Tg treatment, but we saw no evidence of any phosphorylation (data not shown). A more likely explanation is that the phosphorylation status of ATG13 during the Ca^2+^ manipulations signifies whether or not the ULK complex has accumulated in the pre-autophagosomal structures. If the complex has accumulated *and* ULK1 is present, then ATG13 will be phosphorylated, but this phosphorylation is not the driver of the Ca^2+^-dependent response. Because FIP200 and ATG13 are both required for the response, but on their own they cannot mediate it, we suggest that Ca^2+^ affects the assembly (or priming) of those two proteins during the initial stages of autophagy (Fig S6). Once this assembly is underway, ATG9 vesicles will also be contributing to the Ca^2+^ response based on our genetic and imaging data. However, we do not consider that ATG9 vesicles can mediate the response on their own: When we examined FIP200 localization in ATG9 KO MEFs we observed FIP200 puncta upon Tg treatment that were also present without Tg treatment, whereas WIPI2 did not form any puncta (data not shown).

An important corollary of our discovery that, in the absence of ULK1/2, the early autophagy pathway can proceed up to the omegasome step is that the signals that activate the VPS34 complex during autophagy can be independent of mTORC1, ULK1 and AMPK1. One possibility worth exploring is that either ATG9 or the FIP200/ATG13 assemblies can interact with VPS34 directly and help translocate it to the pre-autophagosomal sites. Recent studies have suggested that the protein VMP1 may play a role in lipid transfer and, coupled with the previous observation that VMP1 plays a role in regulating assembly of ER-pre-autophagosomal structures through the dissipation of localised Ca^2+^ signals, provides the interesting possibility that this may underpin the steps missing in this study [47, 48]. Further work is required to confirm this proposal and to understand the upstream cues that may govern the process.

In the process of doing this work, we have examined and experimentally eliminated many potential Ca^2+^ sources as follows: direct effects on ER calcium stores of Bcl-2 or IP_3_R, direct effects on lysosomal stores of TPC1, TPC2, TRPML1, MCOLN1 and MCOLN3, and alterations in mitochondrial Ca^2+^ efflux. Although all of these have been proposed to underlie the Ca^2+^ effect (see Introduction), we could not obtain evidence that they do in our cells. We have also been unsuccessful in showing a clear change in Ca^2+^ dynamics during autophagy, even when using, in addition to the conventional chemical indicators, genetically encoded reporters targeted to the surface of the ER or lysosomes endowing with them with greater sensitivity to detect Ca^2+^ changes at these locations. Again, some evidence has been presented for such dynamics but we could not detect it in our cells. Coupled with the finding that the autophagic response requires only a minimal but sustained Ca^2+^ elevation above baseline, we believe that the source of the Ca^2+^ during autophagy as examined here is a slow localised leak (either from the ER or from an alternative source) that enables the ULK1 complex to nucleate pre-autophagosomal structures in the vicinity of the ER, a hypothesis supported by the finding that BAPTA-2 AM but not EGTA-2 AM was able to inhibit the early autophagosomal structure assembly described here. The requirement for these Ca2+ microdomains also allows for the local Ca2+ signal to be substantially higher than the elevated plateau that we detect through global cellular measurements with Fura-2.

While our manuscript was in preparation, a very recent publication from the Zhang lab reported results of great relevance to those highlighted here. In that work, it was shown that EI24-null cells showed elevated, transient Ca^2+^ signals and that this caused the formation of small, short-lived early autophagosomal structures on the ER surface that required the involvement of FIP200, ATG13 and ATG9 [48]. Interestingly, these calcium signals mediated formation of ULK complex liquid condensates which regulated autophagy initiation. These findings are analogous to the effects shown here – in the case of our work being driven by pharmacological modulation of intracellular Ca^2+^- and provide support for Ca^2+^ being a key signal for autophagosome formation. Whereas Zheng and colleagues proposed various proteins involved in the Ca^2+^ initiation response, further investigation is required to decipher the molecular players that underpin this process including the interplay between ER- and lysosome-derived Ca^2+^ channels.

## Author contributions

NTK designed the project with help from MS and HLR. Experiments were performed by MS, PS, MM, HP. HP and ST provided data on WIPI2 response to Ca^2+^. MS, HLR and NTK wrote the manuscript.

## Acknowledgments

The work was funded by the Biotechnology and Biological Sciences Research Council. We thank Dr Simon Walker for expert help with microscopy, and Dr Martin Bootman for many useful discussions.

