## supplemental data for "Intracellular calcium elevations drive the nucleation of FIP200- and ATG13-containing pre-autophagosomal structures that become omegasomes"

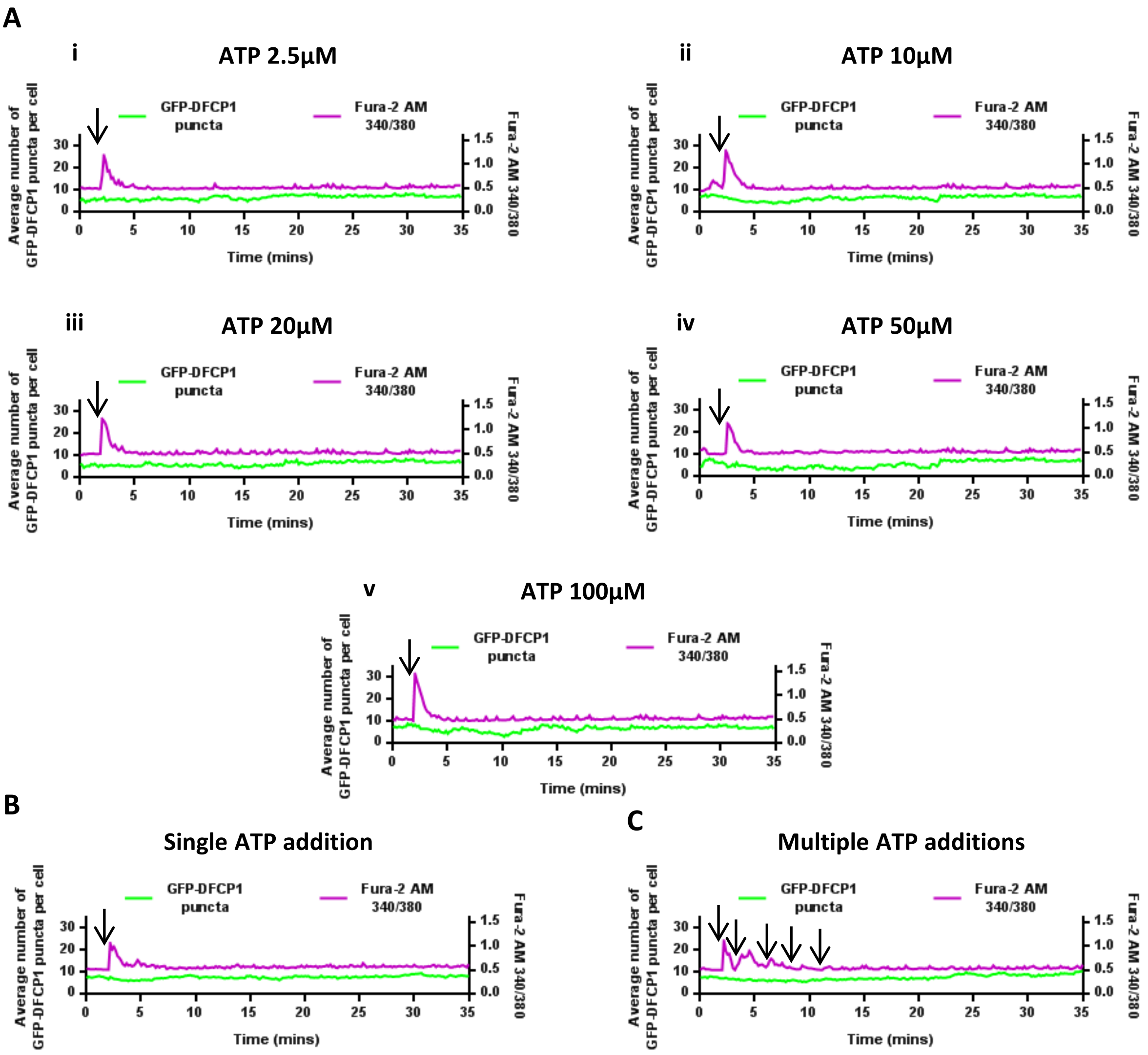


**Supplementary Figure 1.** Enhancing or prolonging the ATP-derived Ca2+ signal does not induce formation of early pre-autophagosomal structures. HEK 293 cells stably expressing GFP-DFCP1 and loaded with Fura-2 AM were imaged live before and after ATP addition (ATP was added 2 min after the start of recording, black arrows). Individual cells were then identified and the GFP-DFCP1 puncta number and Fura-2 AM 340/380 ratio measured in the same cell. (A i-v) ATP was added over a concentration range spanning 2.5 µM and 100 µM. (B,C) Cells set up as above were treated with either a single addition of 2.5 µM ATP (B) or five additions of 2.5 µM ATP at two-min intervals (C). Averages were obtained from analysis of n = 20 cells from two experiments.

Supplementary FIG 1

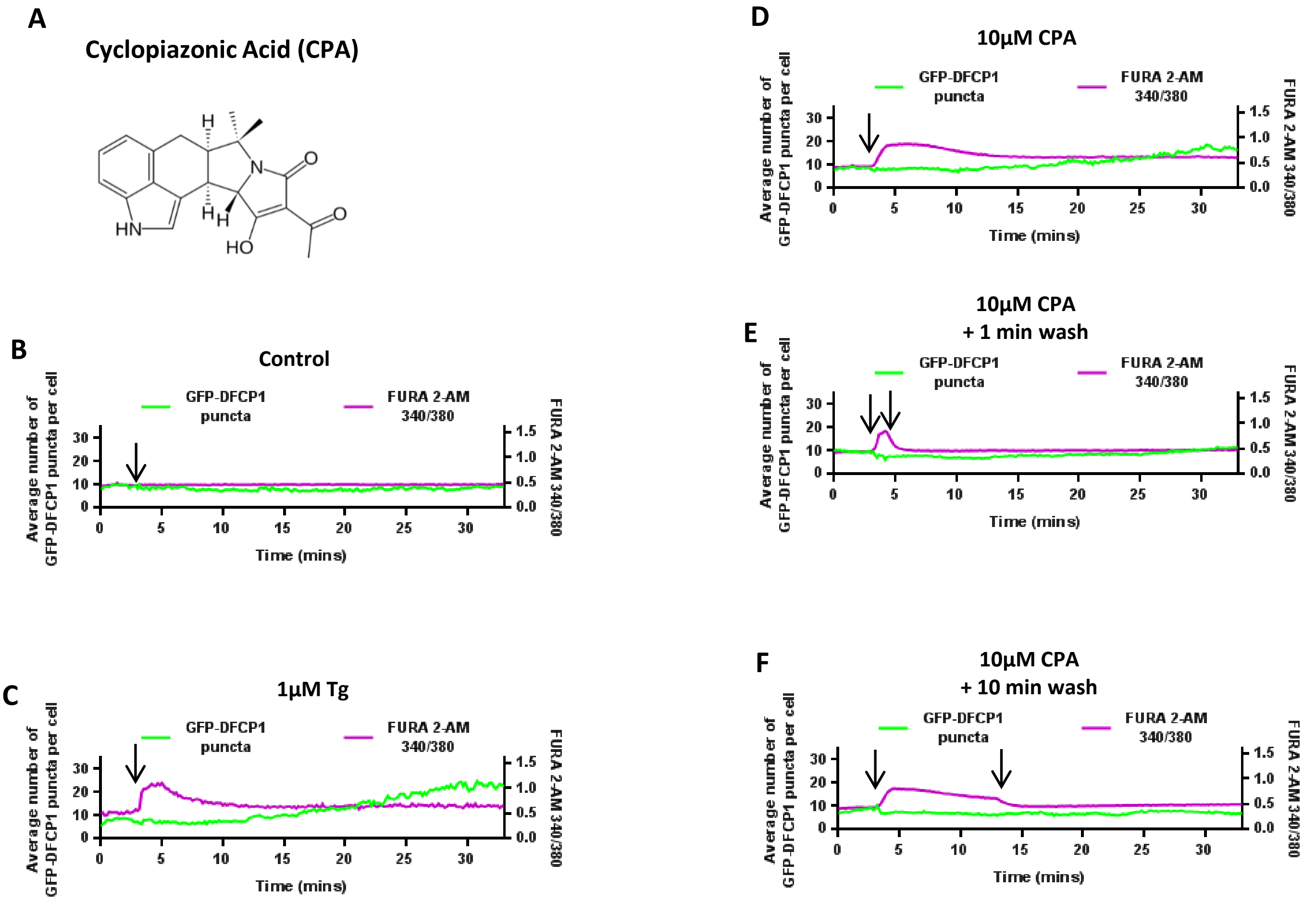


**Supplementary Figure 2.** The reversible SERCA pump inhibitor cyclopiazonic acid increases GFP-DFCP1 puncta in fed conditions but not if washed out after either 1 or 10 min of addition. HEK 293 cells stably expressing GFP-DFCP1 were loaded with Fura-2 AM and imaged live. Individual cells were then identified and the GFP-DFCP1 puncta number and Fura-2 AM 340/380 ratio measured in the same cell. (A) Chemical structure of CPA. Cells were imaged for 3 min to obtain a baseline after which they were treated with either vehicle (DMSO) (B), 1 µM Tg (C) or 10 µM CPA (D) 3 min (black arrows). In panels E and F, the cells were washed with control medium at the time point indicated by the second black arrow.

Supplementary FIG 2

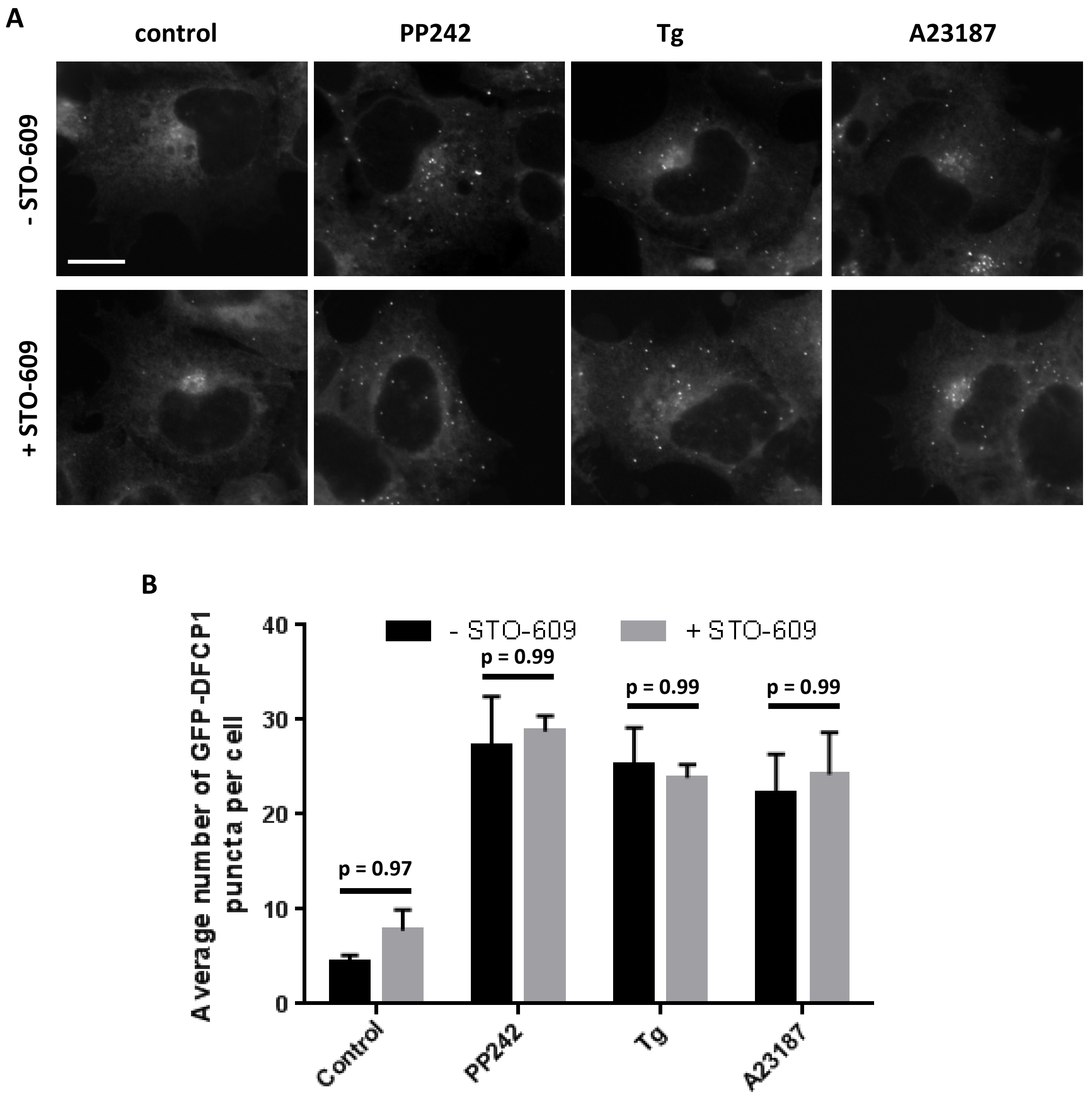


**Supplementary Figure 3.** Tg and A23187 do not activate autophagy through AMPK signalling. HEK 293 cells stably expressing GFP-DFCP1 were treated with 1 µM PP242 (1 h), 1 µM Tg or 10 µM A23187 (30 min) with or without 50 µM STO-609. After fixing, the samples were stained for WIPI2 (A) and average puncta per cell were plotted (B). (n = 3, ± SEM). Scale bar = 10 µm.

Supplementary FIG 3

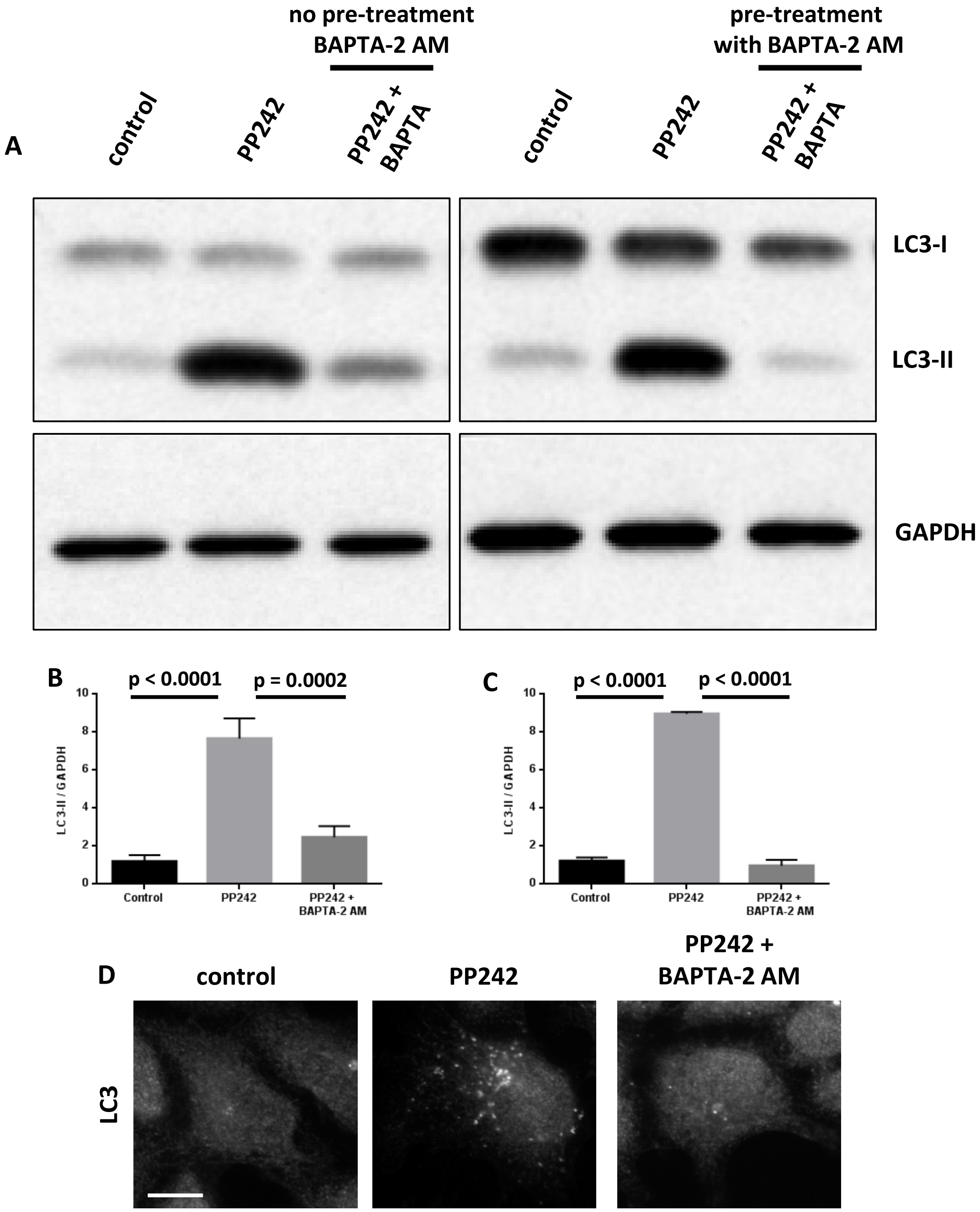


**Supplementary Figure 4.** Cell loading with BAPTA-2 AM inhibits formation of LC3-positive autophagosomes. (A) HEK 293 cells were treated with 1 µM PP242 or PP242 in the presence of 10 µM BAPTA-2 AM. BAPTA-2 AM was either added simultaneously with PP242 (left panels) or 5 min before PP242 (right panels). Samples were lysed and LC3 levels were examined by immunoblotting. Quantitation of these experiments is shown in panels B and C. (D) HEK 293 cells were treated with vehicle, 1 µM PP242 or PP242 in the presence of 10 µM BAPTA-2 AM. Cells were fixed and immunostained for endogenous LC3 levels. Scale bar approximately 10 µm. (n = 3, ± SEM).

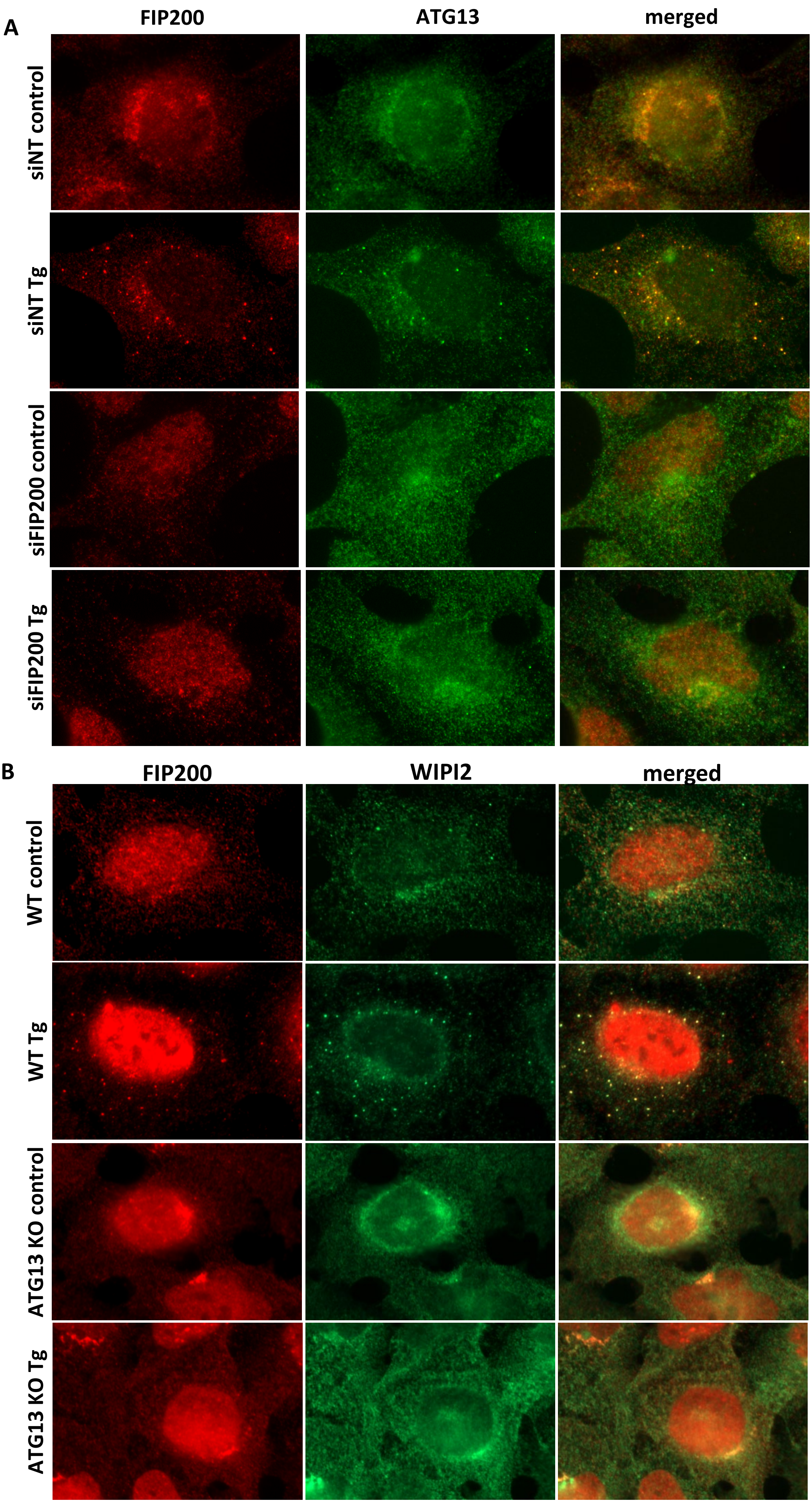


Supplementary FIG 5

**Supplementary Figure 5.** FIP200 and ATG13 are together required for the calcium-induced response. (A) HEK 293 cells in which FIP200 expression was reduced by siRNA transfection and subsequent culture for 72 h were treated with 1 µM Tg for 30 min. Following fixation, cells were immunostained for endogenous ATG13 and FIP200. Note the reduction of FIP200 signal which indicates good siRNA down-regulation. (B) HEK 293 cells with a CRISPR-mediated deletion for ATG13 or parental HEK 293 cells were treated with 1 µM Tg for 30 min and immunostained as above for endogenous FIP200 and WIPI2.

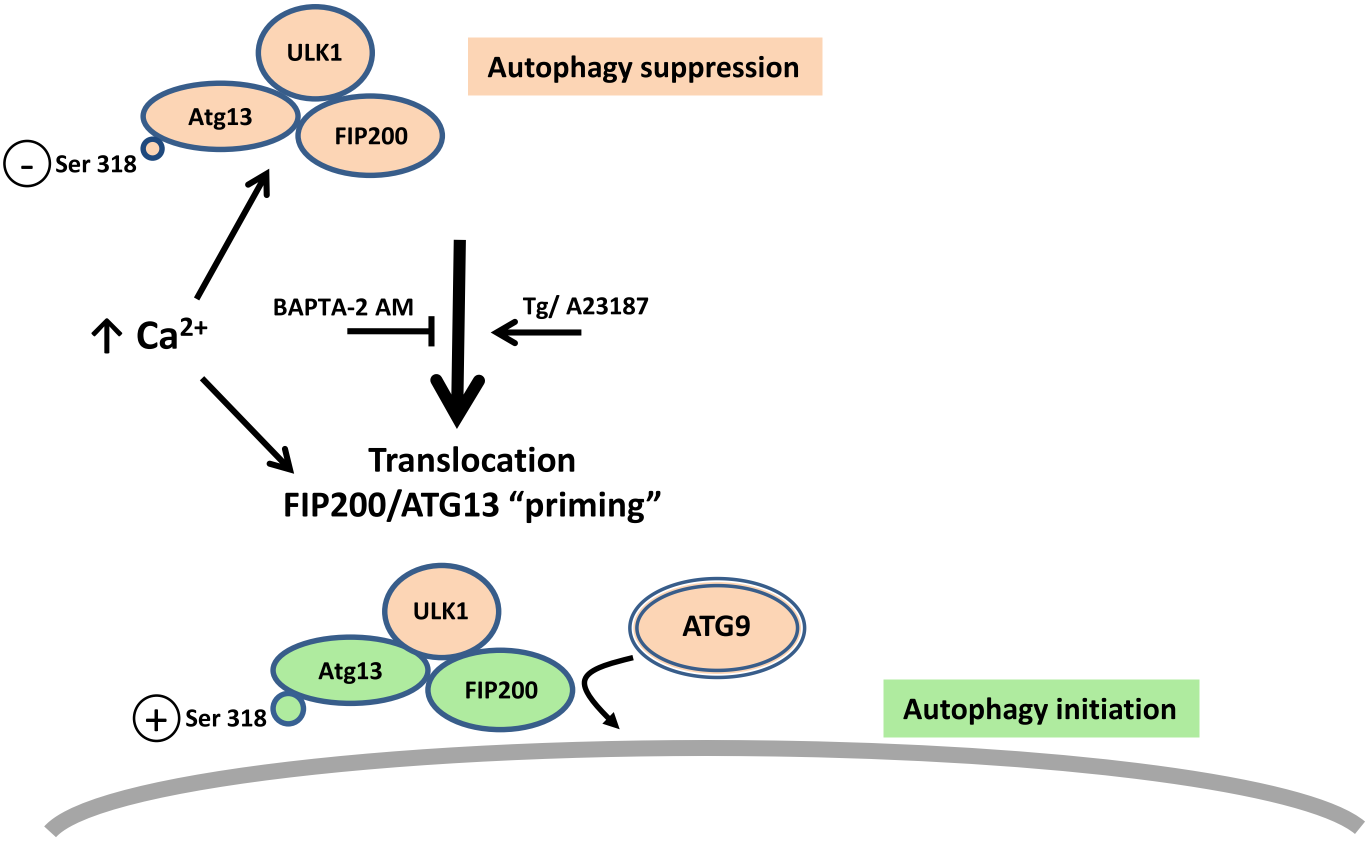


**Supplementary Figure 6.** Effect of cytosolic Ca2+ levels on ULK complex during early autophagy. When cytosolic Ca2+ levels are elevated for a prolonged duration by Tg or A23187, the ULK complex translocates to the site of autophagosome initiation (frequently on the ER). This translocation can take place in complete medium, independently of mTORC1 status and in the absence of ULK1 protein. It does however require both FIP200 and ATG13. Upon mTORC1 inactivation and in the process of autophagy induction, chelation of cytosolic Ca2+ with BAPTA-2 AM inhibits autophagy, including the enhanced phosphorylation of ATG13 at Ser318. Under these conditions, the ULK complex does not translocate and autophagy is not initiated.
